# Social and environmental cues mask circadian activity patterns in a highly social shell-dwelling cichlid fish

**DOI:** 10.64898/2026.08.07.741809

**Authors:** Laura Fritschi, Annika L. A. Nichols, Rita Gonzalez-Dominguez, Adrian Indermaur, Attila Rüegg, Walter Salzburger, Maxwell E.R. Shafer

## Abstract

Species exhibit variation in their circadian activity rhythms, including being active during the day (diurnal), the night (nocturnal), or the twilight periods (crepuscular). While the influence of environmental factors such as light on entraining and controlling activity patterns is well known, it is unclear how social and abiotic factors affect circadian behaviors. The daily activity patterns of Lake Tanganyikan cichlid fish are diverse, and may be associated with species diversification. Intriguingly, some shell-dwelling cichlid species, which anecdotal observation suggests are diurnal, displayed strong nocturnal activity when assayed in a “common garden” lab setup. Here, by integrating field and lab-based studies, we provide three lines of evidence that social and environmental cues mask underlying circadian rhythms in shell-dwelling cichlids. First, we demonstrate that shell-dwelling cichlids are diurnal in their natural habitat, but convert to nocturnal activity when assayed alone in a common garden. Second, we identify that in the presence of a shell and conspecifics the highly social *N. multifasciatus* becomes diurnal. In contrast, the circadian activity pattern of a closely related, but sub-social species, *N. brevis*, is masked only by a shell, and unaffected by the presence of conspecifics. Third, we demonstrate that the masking effect of shells and conspecifics is circadian and continues in the absence of light. Fourth, we identify that the response to these cues is influenced by sex, with greater effects in female fish, and by pedigree, with stronger effects in individuals bred and raised in captivity compared to those captured in the wild. Together, these experiments reveal new relationships between circadian rhythms and sociality in these fishes, offering broader insights into the ecological and evolutionary drivers of these behaviours.

## Introduction

Every animal species exhibits characteristic daily activity patterns, or circadian rhythms, such as nocturnal (active at night), diurnal (active during the day), crepuscular (active at dawn and dusk), or cathemeral (showing irregular or arrhythmic activity across the 24-hour cycle). These endogenous circadian rhythms can be influenced by different biotic and abiotic factors (Vaze and Sharma, 2013), such as daily light cycles, temperature, activity patterns of competitors and/or predators (Acerbi and Nunn, 2011; Fenn and Macdonald, 1995; Lear et al., 2021; Levy et al., 2007; Nguyen et al., 2020; Siegel, 2009), food availability, and parental care (Reebs and Colgan, 1992, 1991). However, little is known about the roles of within-species social factors and the interplay of environmental cues other than light on the regulation of temporal activity patterns.

Temporal activity patterns are regulated by the endogenous circadian clock (process C)^6,9^. Process C describes a molecular mechanism that synchronises an inner clock gene cascade to the outer world through entrainment factors. These factors act as reference points throughout the day and create the iterative 24h activity pattern. Increases in ambient light in the morning is the most powerful entrainment factor for process C (Borbély et al., 2016; Halle and Stenseth, 2000), but other external stimuli simultaneously synchronise the underlying rhythm such as food availability (Beale et al., 2013; Moran et al., 2014), or the presence or activity of conspecifics (Castillo-Ruiz et al., 2012). There can be additional players downstream of process C, known as masking factors. Masking factors differ from entrainment factors, because they do not change the underlying clock (process C), but can change the output of the clock once they appear (Mrosovsky, 1999).

Across the animal kingdom, many species display phase shifts in activity driven by various masking factors. A striking example of this is the plastic response of the golden spiny mouse (*Acomys russatus)* (Levy et al., 2007), which displays a diurnal activity pattern in the wild, but switches to nocturnal activity when transferred to the lab (Levy et al., 2007). Geckos in the genus *Cnemaspis* have an endogenous diurnal activity pattern^4^, which is masked by the presence of interspecific competition from species of the genus *Cyrtodactylus*. A switch in phase of temporal activity can also be triggered by an antipredatory response. For example, a subpopulation of Norway rats (*Rattus norvegicus*) can shift their temporal activity pattern to avoid predation by red foxes (*Vulpes vulpes*) (Fenn and Macdonald, 1995). The mutual effect and selective pressures of antagonistic species on activity patterns is relatively well understood. Far less is known about how conspecifics and complex social interactions influence daily activity patterns, though group-living species often sleep collectively rather than in isolation (Chakravarty et al., 2024). Social interaction and conspecific activity patterns can entrain the endogenous clock in honey bees (Fuchikawa et al., 2016; Siehler et al., 2021), and social housing or interactions are potent regulators of activity, and are often associated with convergent activity rhythms in multiple species (Castillo-Ruiz et al., 2012). However, it is difficult to disentangle both the entrainment and masking effects of social interactions, and the effects of other environmental cues.

Disentangling the factors that shape a species’ diel activity patterns requires an experimental model that displays variation in activity in response to such cues. Cichlid fishes are an established model system for studying the evolutionary and ecological correlates of behavior (Maruska and Fernald, 2018). Cichlids from Lake Tanganyika show unparalleled variation in morphological, ecological, and behavioral phenotypes, allowing them to inhabit a wide range of ecological niches (Ronco et al., 2021, 2020; Salzburger, 2018; Salzburger et al., 2014; Takahashi and Koblmüller, 2011). Cichlids from Lake Tanganyika (Nichols et al., 2025) and Lake Malawi (Lloyd et al., 2024, 2021) have also diversified along the temporal axis leading to nocturnal, diurnal, crepuscular, and cathemeral species. Shell-dwelling cichlids from Lake Tanganyika are the smallest cichlid species and exhibit a sedentary lifestyle, specialising in the use of empty *Neothauma tanganyicense* shells for shelter and breeding (Arnegard and Carlson, 2005; Frøland Steindal and Whitmore, 2019; Kohler, 1998; Loftus et al., 2022; Mcglue et al., 2010; Peart et al., 2014; Rossiter, 1995). Shell-dwelling cichlid species display different levels of social complexity, ranging from the highly social *N. multifasciatus* to the solitary living *N. brevis* (Kohler, 1998; Jordan et al., 2021, 2016; Gübel et al., 2021; Sato and Gashagaza, 1997; Schradin and Lamprecht, 2002; Jordan et al., 2026). Though anecdotal observations have suggested that these species are diurnal in the wild, they were recently identified as having strictly nocturnal activity patterns in the lab when assayed in a “common garden” setup (Nichols et al., 2025), suggesting that their activity patterns could be affected by masking factors.

In this study, we demonstrate that in their natural habitat, multiple species of shell-dwelling cichlids show clear diurnal activity patterns, but when observed in a common garden experiment in the laboratory, they switch to nocturnal activity patterns. We identify that in *N. multifasciatus*, both the presence of shells and conspecifics are required to maintain diurnal activity patterns under laboratory conditions, and that this effect is further modulated by the sex and genetic background of individuals. In contrast, in the less social *N. brevis*, only the presence of shells, but not conspecifics, are necessary for diurnal activity. Together these experiments demonstrate how species-specific activity patterns are influenced by life-history, and social and environmental cues, revealing a novel relationship between the circadian timing of sleep and sociality.

## Results

### Shell-dwelling cichlid species display diurnal activity patterns in the wild

Previous observations of multiple species of shell-dwelling cichlids in a laboratory setting suggest that they have strong nocturnal activity patterns (Nichols et al., 2025). However, anecdotal reports from *in situ* observations suggest that shell-dwelling cichlids are diurnal in their natural environment (*personal communication, A. Indermaur*). To examine their baseline behavior in the wild, we recorded groups of three cichlid species in Lake Tanganyika (*Neolamprologus multifasciatus, Lamprologus ocellatus,* and *Telmatochromis temporalis)* that were included in our previous study (Nichols et al., 2025) (**Supplemental videos 1-3**). Fish were recorded in 10 min intervals over 24 hour periods. *N. multifasciatus* were observed in large colonies, consistently interacting with conspecifics, whereas the *L. ocellatus* and *T. temporalis* individuals were solitary, inhabiting isolated shells on the sandy substrate. Quantification of actively moving and visible individuals from field recordings strongly support diurnal activity patterns of all species examined (**Figure 1** and **Supplemental Data 1**). Individuals of each species consistently displayed the same activity pattern across all recorded days. During the daytime, fish were frequently observed defending territories, feeding, and spitting sand into neighboring territories. At night, fish were rarely visible and, when observed, were typically located in or near their shells (**Figure 1C-K**). Lake Tanganyika harbors nocturnal piscivorous catfish, and we recorded multiple nighttime attacks on our focal cichlids. In all cases, the cichlids successfully escaped into their shells (**Supplementary video 1**). These observations demonstrate that, in their natural habitat, shell-dwelling cichlids are diurnal.

**Figure 1:**
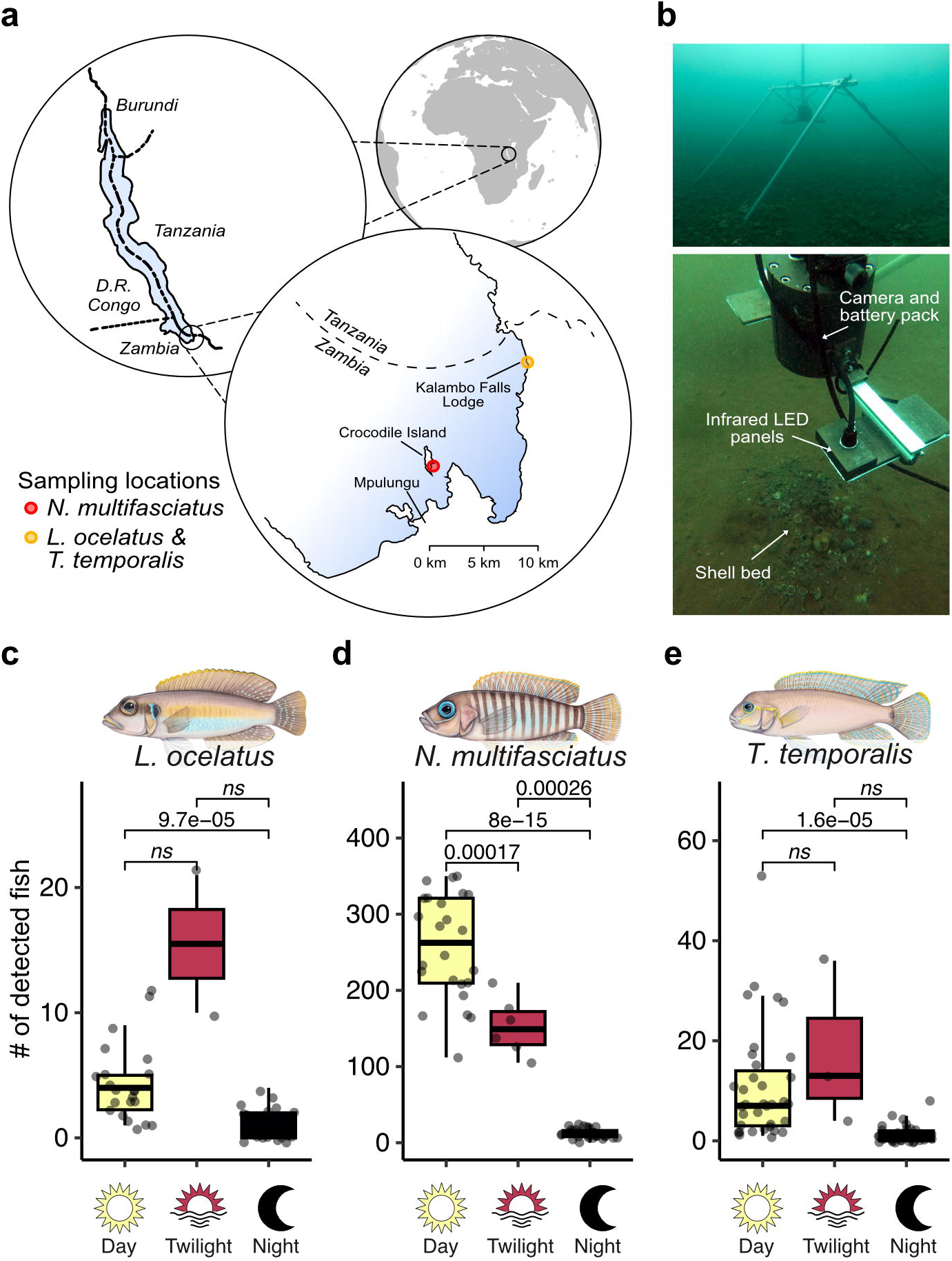
Shell-dwelling species of cichlids are diurnal in the wild. **(a**) The study location in Lake Tanganyika. Shell dwelling cichlids were filmed *in situ* in two locations in Zambia on the Southern coast of Lake Tanganyika. (**b**) Field setup noting the position of the tripod, infrared lights, and infrared camera. *Photograph by Adrian Indermaur.* Quantification of the number of fish movements (in and out of frame) *L. ocellatus* (**c**), *N. multifasciatus* (**d**), and *T. temporalis* (**e**) across the diel cycle. Boxplots show the interquartile range (limits of boxes correspond to 1st and 3rd quartiles), whiskers represent 1.5*(interquartile range), and the line is the mean.

### Captive and wild caught populations display nocturnal activity patterns in a laboratory setting

Shell-dwelling cichlids displayed diurnal activity patterns in the wild, but nocturnal activity patterns in a reductionist assay in captivity (Nichols et al., 2025). These results suggest that either laboratory rearing or breeding affects activity patterns, or that previous experiments lacked specific environmental cues necessary to elicit their natural diurnal activity. In this study, we focused on *N. multifasciatus,* as it is readily available and easy to house in a laboratory setting. To test whether pedigree and laboratory rearing affects their diel activity, we compared three different populations of *N. multifasciatus* (an aquarium strain: AS; a laboratory strain: LS; a Zambian population: ZP), which represent variations in how they were raised and the number generations they have spent in captivity (**Figure 2a**). The AS has been maintained in captivity by commercial breeders for numerous generations. In contrast, the LS has only experienced a few generations in captivity, while the ZP was collected from Africa shortly before the experiment and had no prior history of captivity. Activity was measured by calculating the average movement speed of each individual across the diel cycle for six days and six nights (**Figure 2b**) (Nichols et al., 2025).

**Figure 2:**
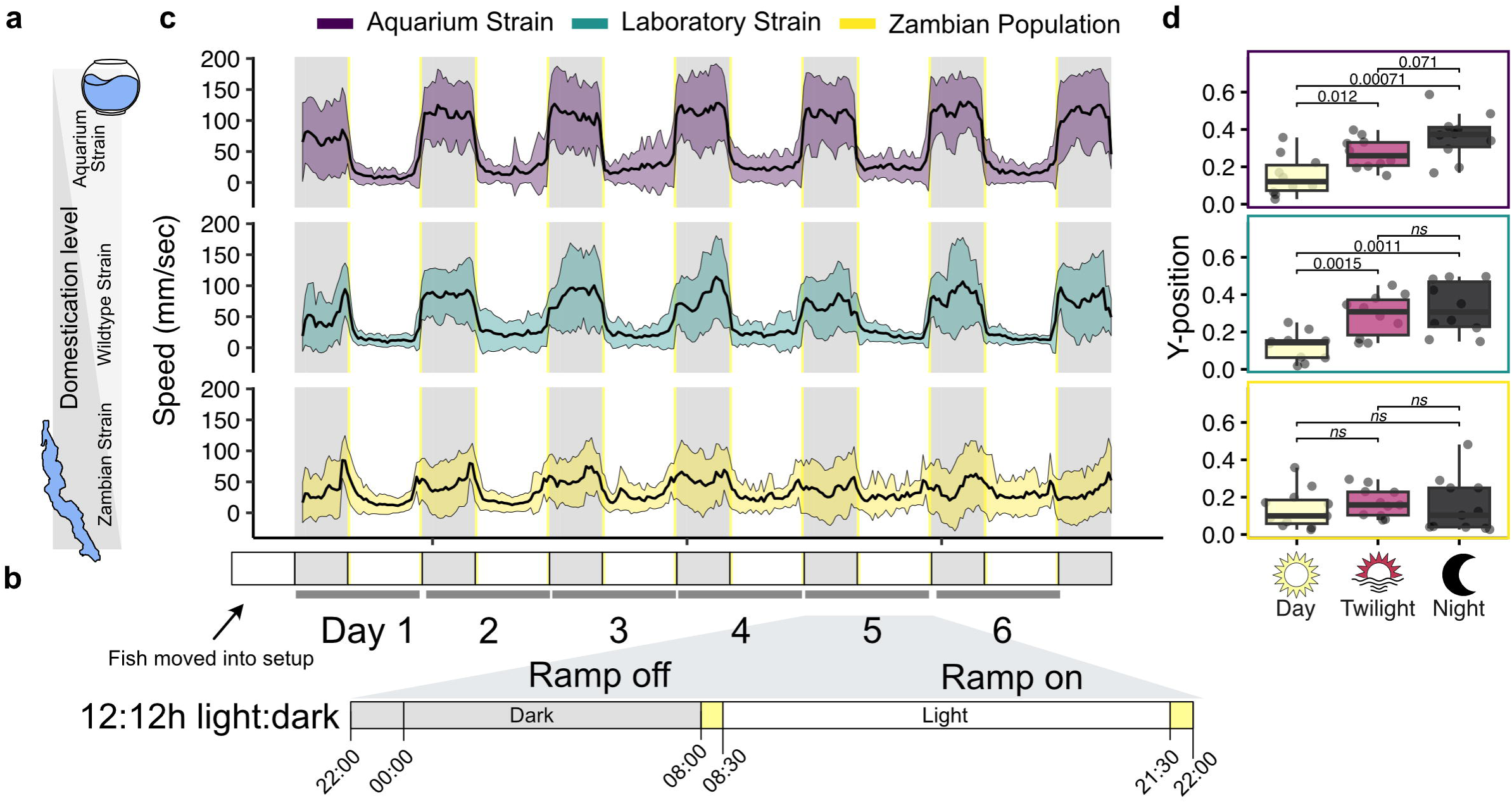
*Neolamprologus multifasciatus* strains display a nocturnal activity pattern in captivity regardless of pedigree. (**a**) *N. multifasciatus* strains represent varying degrees of domestication, from the fully domestic “Aquarium Strain”, lab bred “Laboratory Strain”, and wild-type “Zambian Population”. (**b**) Schematic of light cycle for in lab tracking of activity patterns. Grey areas indicate night when lights are off, yellow areas indicate dawn and dusk when lights are ramping up or down, and white indicate daytimes when lights are on. (**c**) Activity pattern of each *N. multifasciatus* strain averaged across 30 minute bins, and across individuals over 6 days and 6 nights. (**d**) Quantification of the average relative y-position (height in tank) during the day, twilight (dawn and dusk), and night, for each of the three strains of *N. multifasciatus*. Boxplots show the interquartile range (limits of boxes correspond to 1st and 3rd quartiles), whiskers represent 1.5*(interquartile range), and the line is the mean.

Overall, we observed that all three populations displayed increased activity at night across six consecutive days and nights (**Figure 2c**). The AS expressed the highest degree of nocturnal activity, followed by the LS, whereas the ZP displayed the lowest activity at night (ANOVA *p-value* = 0.00257). In our field recordings, these cichlid species hovered above their shells during the day, and rested near their entrance during the night (**Supplementary Video 1**). However, in the laboratory, the AS and LS strains swam significantly higher in the tank during the night than the day, associated with their fastest swimming speeds and highest activity levels (**Figure 2d**). Our results suggest that the nocturnal laboratory behavior is observed across aquarium and wild-caught strains, with significantly increased night-time activity in strains raised in captivity. Nonetheless these experiments indicate that a specific environmental cue or cues present in the wild, but absent in the laboratory, rather than the effect of domestication or laboratory housing, drives the shift from diurnal to nocturnal behavior.

### A combined set of environmental factors are required for diurnal activity in *N. multifasciatus*

All three strains of *N. multifasciatus* had nocturnal activity patterns in the lab, including individuals that were held in captivity for less than 2 months. In our experimental setups, fish were housed in tanks with only a sandy substrate and no other enrichment, and were unable to interact with conspecifics. Importantly, their natural habitat in Lake Tanganyika consists of numerous empty shells (*Neothauma tanganyicense*) that have been brought together through water movements to form massive shell beds (Bose et al., 2020a; Busch et al., 2018; Cohen and Thouin, 1987; Hori et al., 1993; Mcglue et al., 2010). Shell-dwelling cichlids congregate on these shells for both shelter and breeding, and display strong site fidelity and social interactions. We therefore hypothesised that the endogenous activity patterns of shell-dwelling cichlids may be masked by the absence of these shells or through the effect of social isolation. To test whether shells or the lack of conspecifics were masking factors, we examined each of the three strains of *N. multifasciatus* under four different conditions. We tested their diel activity in absence of all cues, in the presence of a shell, in the presence of conspecifics, or in the presence of both shells and conspecifics using the same assay as previously (**Figure 2b**, and **Supplemental data 2-3**). For conspecific conditions, the examined species were separated into groups of three one week prior to the experiment to establish familiarity. During recordings, one focal fish was placed in an experimental arena and the remaining two individuals were placed in an adjacent arena, separated by a mesh divider that allowed for visual and olfactory interactions. All individuals were tracked as previously described.

Overall, both shells and conspecifics decreased nocturnal activity, but when fish were exposed to them in combination, produced a crepuscular activity pattern for both the AS and LS (**Figure 3a**). The combination of cues also caused a decrease in average vertical position during the twilight and night in both the AS and LS (**Figure 3b**). In contrast, the presence of the cues had a weaker, and non-significant, effect on the ZP. These results suggest that both shells and conspecifics can act as masking factors in domesticated strains, and that the absence of additional cues might still be affecting the behaviour of the Zambian strain.

**Figure 3:**
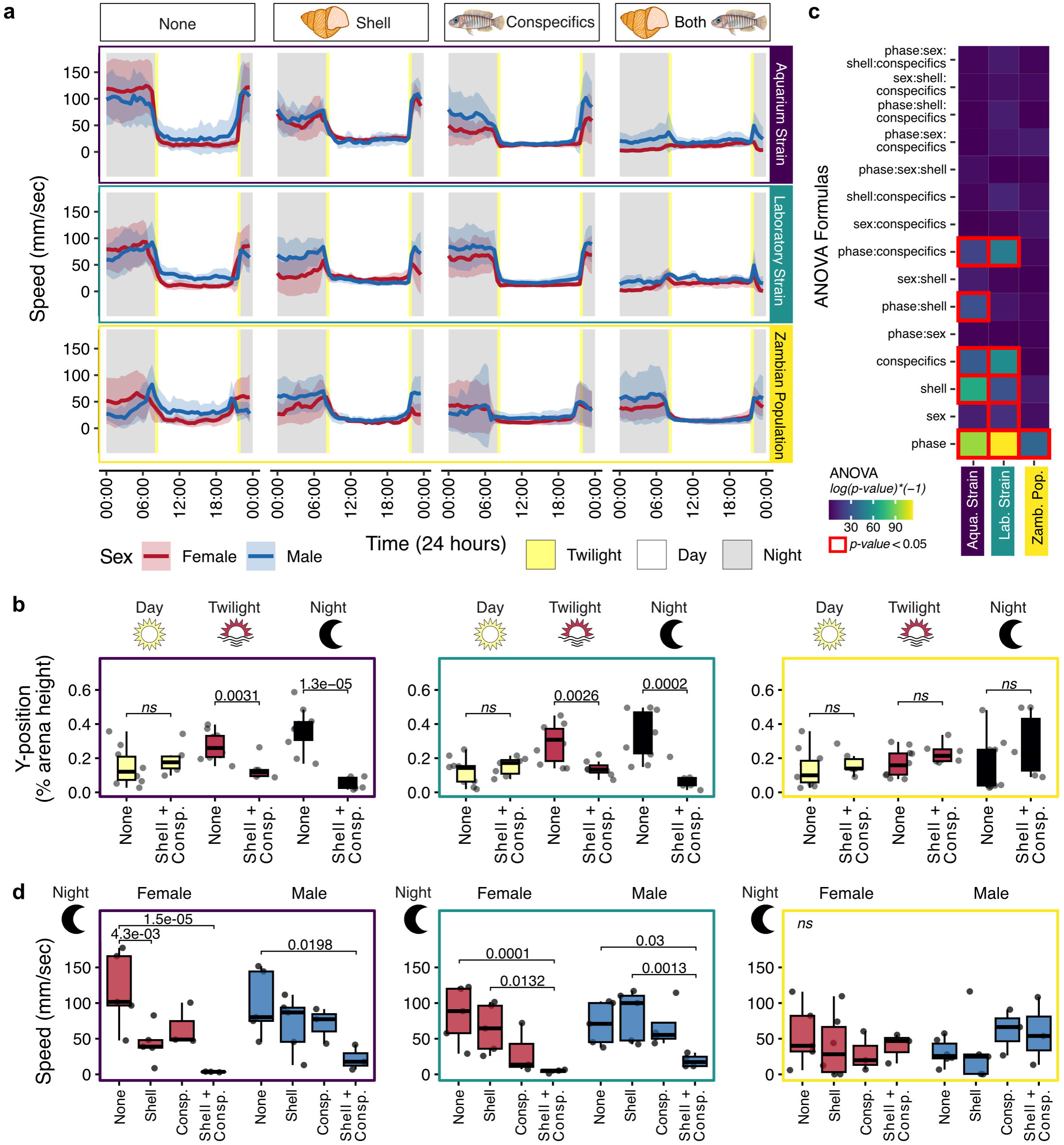
The contribution of each masking factor is affected by sex and pedigree. (**a**) Plots showing the average daily activity patterns for *N. multifasciatus* in the absence of cues, presence of a single shell, presence of only conspecifics, or in the presence of both a shell and conspecifics. Data are separated by strain, and the average activity patterns of both female and male fish are displayed. Each plot shows the activity patterns of 10-12 individuals, averaged across days. Shaded ribbons display 1 standard deviation from the mean. Grey shaded areas indicate nighttime, yellow is dawn and dusk, and white areas are daytime. (**b**) Results of an ANOVA for the interaction between phase (day, twilight, night), sex (male or female), presence of a shell (yes or no), and presence of conspecifics (yes or no) on the swimming speed of fish from each strain of *N. multifasciatus*. (**c**) Quantification of the average relative y-position (height in tank) during the day, twilight (dawn and dusk), and night, for each of the three strains of *N. multifasciatus.* (**d**) Quantification of the average swimming speed during the night, for female and male fish from each of the three strains of *N. multifasciatus.* Boxplots show the interquartile range (limits of boxes correspond to 1st and 3rd quartiles), whiskers represent 1.5*(interquartile range), and the line is the mean.

All three strains also appeared to have sex-specific differences in their response to either or both cues suggesting a complicated relationship between sex, domestication, and each cue. To more systematically test for combinatorial effects, we analyzed phase specific averages for speed using Analysis of Variance (ANOVA) accounting for presence of environmental cues (shells and conspecifics), strain, and sex, as well as their interactions, followed by post-hoc Tukey’s HSD tests (**Figure 3c**). This analysis revealed that the activity patterns of all three *N. multifasciatus* strains were described by phase, sex, and the presence of either or both cues, as well as combinatorial effects of some of the variables (**Figure 3c**). In particular, the nocturnal activity of females of the AS and LS strains were significantly reduced by the combination of cues, with individual cues causing intermediate effects. In contrast, individual cues had weaker effects on males of those strains, whose nocturnal activity was only significantly reduced by the presence of both cues in combination (**Figure 3d**). Males and females of the ZP strain deviated from this pattern, remaining nocturnal under all four conditions (**Figure 3c-d**). These experiments demonstrate that the contributions of shells and conspecifics to the activity patterns of *N. multifasciatus* is affected by both sex and pedigree.

### Species sociality level changes the impact of the masking factors

The results from our experiments with *N. multifasciatus* suggest that all three strains are nocturnal in a reductionist assay, but differ in their set of environmental and social factors to maintain diurnality in the lab. Importantly, different species of shell-dwelling cichlids display variations in their social complexity (Jordan et al., 2021; Lein and Jordan, 2021). To test whether a species’ social complexity affects their reactions to masking factors, we examined *Neolamprologus brevis*, a closely related species of *N. multifasciatus*. *N. brevis* occupy a similar ecological niche in Lake Tanganyika and are about the same size, but they are less social than *N. multifasciatus* (Fryer and Iles, 1972; Jordan et al., 2016). They live in solitary breeding pairs and protect their home shell from all potential intruders. We hypothesised that, in contrast to *N. multifasciatus*, shells, but not conspecifics, would act as masking factors in *N. brevis*. To test the impact of shells and conspecifics on their diel activity, we recorded individuals of *N. brevis* under the same four conditions used for *N. multifasciatus* (no cues, only a shell, only conspecifics, or both a shell and conspecifics). As before we performed ANOVA to determine which cues or combinations of cues affected phase-specific activity patterns.

We observed that the activity patterns of *N. brevis* were described by the presence or absence of a shell in combination with phase, but not the presence of conspecifics (**Figure 4a**). Indeed, individuals of *N. brevis* switched from a nocturnal pattern to a diurnal pattern in response to a shell only, whether or not it was in combination with conspecifics (**Figure 4b**). The presence of the shell caused a significant decrease in activity (swimming speed) as well as a reduction in average y-position during both the twilight and night, regardless of sex (**Figure 4c**). No effect of conspecifics was observed, and both sexes of *N. brevis* were nocturnal in the absence of either cue, but generally less active at night compared to *N. multifasciatus* (**Figure 3a**). Together, these experiments illustrate that the contribution of each of these environmental factors is not only affected by sex and pedigree, but also by the sociality and life history traits of a species.

**Figure 4:**
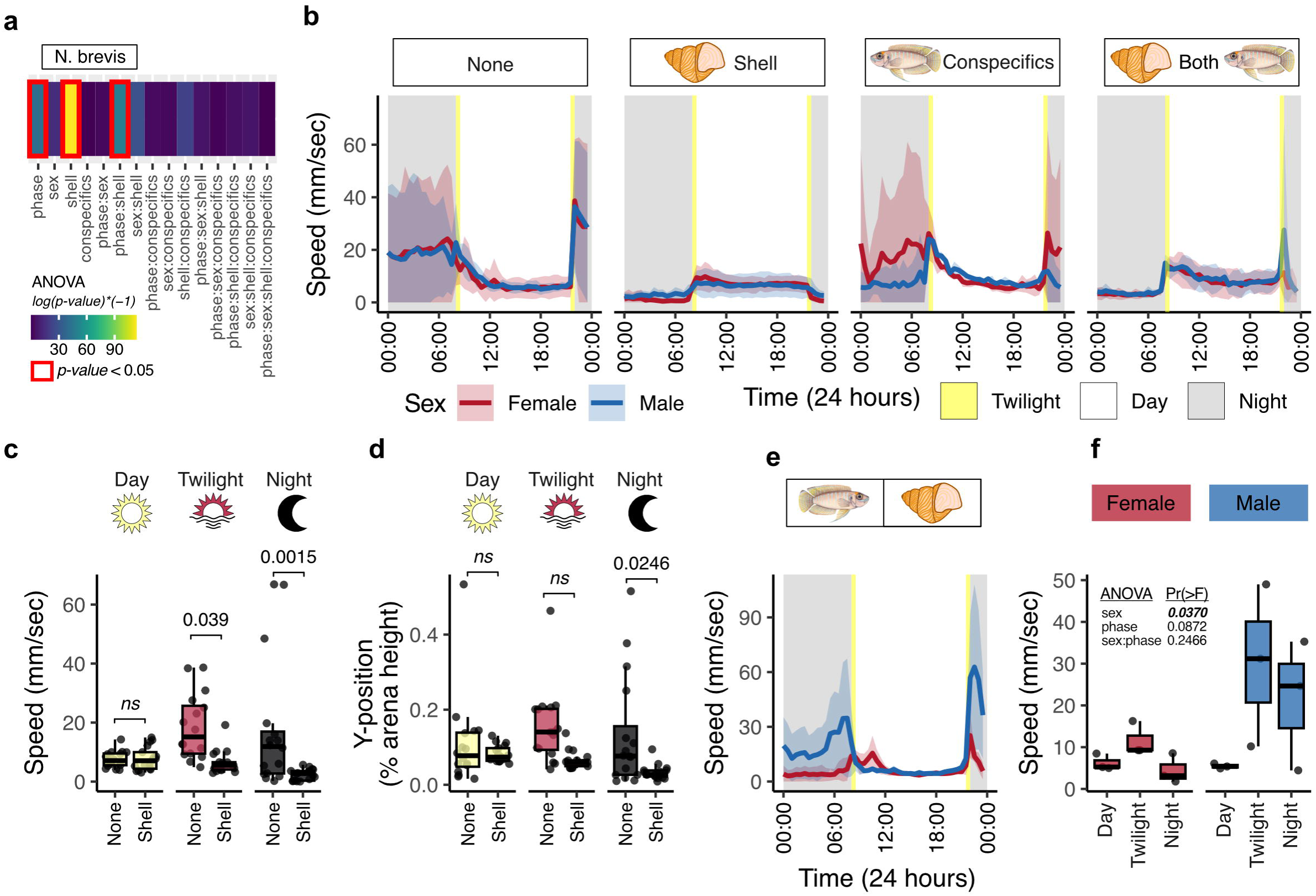
*Neolamprologus brevis* are diurnal in the presence of a shell. (**a**) Results of an ANOVA for the interaction between phase, sex, presence of a shell, and presence of conspecifics on the swimming speed of fish from each strain of *N. brevis*. (**b**) Plots showing the average daily activity patterns for *N. brevis* in the absence of cues, presence of a single shell, presence of only conspecifics, or in the presence of both a shell and conspecifics. The average activity patterns of both female and male fish are displayed. Each plot shows the activity patterns of 10-12 individuals, averaged across days. Shaded ribbons display 1 standard deviation from the mean. Grey shaded areas indicate nighttime, yellow is dawn and dusk, and white areas are daytime. (**c**) Quantification of the average swimming speed during the day, twilight (dawn and dusk), and night, for *N. brevis.* (**d**) Quantification of the average relative y-position (height in tank) during the day, twilight (dawn and dusk), and night, for *N. brevis.* (**e**) Plot showing the average daily activity patterns for *N. brevis* in the absence of cues, but with a visible shell in the adjacent arena. The average activity patterns of both female and male fish are displayed. Each plot shows the activity patterns of 10-12 individuals, averaged across days. Shaded ribbons display 1 standard deviation from the mean. Grey shaded areas indicate nighttime, yellow is dawn and dusk, and white areas are daytime. (**f**) Quantification of average speed during the day, twilight (dawn and dusk), and night for male and female fish exposed to no cues but a visible shell in the adjacent arena. Boxplots show the interquartile range (limits of boxes correspond to 1st and 3rd quartiles), whiskers represent 1.5*(interquartile range), and the line is the mean.

Though we did not observe a significant effect of conspecifics alone on the activity patterns of individual *N. brevis*, nocturnal activity under these conditions was slightly depressed compared to the no cue condition (**Figure 4b**). Given the strength of the effect of shells in modulating the activity patterns of *N. brevis*, we wondered if the shells in the adjacent arena used by the conspecifics were affecting the activity of the focal fish, who could see shells, but were unable to directly interact with them. To test whether or not the visual cue of a shell alone could modulate the circadian activity of *N. brevis* individuals, we placed an individual shell in the adjacent arena and recorded the activity of individual *N. brevis*. Surprisingly, this experiment revealed that a visual shell cue was sufficient to shift the activity pattern in females, but only a physical shell could mask activity patterns in males (**Figure 4e-f**). These results suggest that visual cues alone can act as masking factors in shell-dwelling cichlids, but their effect may be sex-specific.

### Masked and unmasked behaviors are circadianly regulated in *N. multifasciatus*

In fish and other animals, light can act as both an entrainment cue, synchronizing the internal circadian clock with the external environment, and as a masking factor that enhances or suppresses activity patterns. Given that the observed activity patterns occur immediately, and without ‘jet-lag’, our results suggest that shells and conspecifics are masking factors that potentially reverse the phase of the activity patterns of shell-dwelling cichlids. However, an interaction between the masking effect of light with the effects of shells and conspecifics could also explain our results, such that in the absence of all cues, locomotor activity is enhanced under dark nocturnal conditions. To disentangle the masking effects of light and social and environmental cues, we performed light cycle manipulations in the presence and absence of shells and conspecifics. We exposed each fish to 48 hrs of normal light:dark cycle, followed by 2 dark and 2 light pulses during the next 72 hrs, followed by 72 hrs of a dark:dark cycle, and finally 72 hrs of normal light:dark cycle.

First, we investigated the activity patterns of *N. multifasciatus* in response to complete darkness (dark:dark), followed by their response to dark pulses during the day and light pulses during the night (**Figure 5**). Individual fish were recorded without masking factors or in presence of a shell and conspecifics. Under dark:dark conditions, both sexes maintained the same activity pattern as under light:dark, adopting a ‘nocturnal’ activity pattern without shells and conspecifics and a diurnal/crepuscular activity pattern in presence of a shell and conspecifics (**Figure 5a-b**). The activity patterns (swimming speeds) for fish with and without masking factors, and during both light:dark and dark:dark epochs also had periodogram peaks around 24 hrs (**Figure 5c**). This result confirms that both diurnal and nocturnal activity patterns are circadian, and that shells and conspecifics act as masking factors to reverse the endogenous circadian activity pattern.

**Figure 5:**
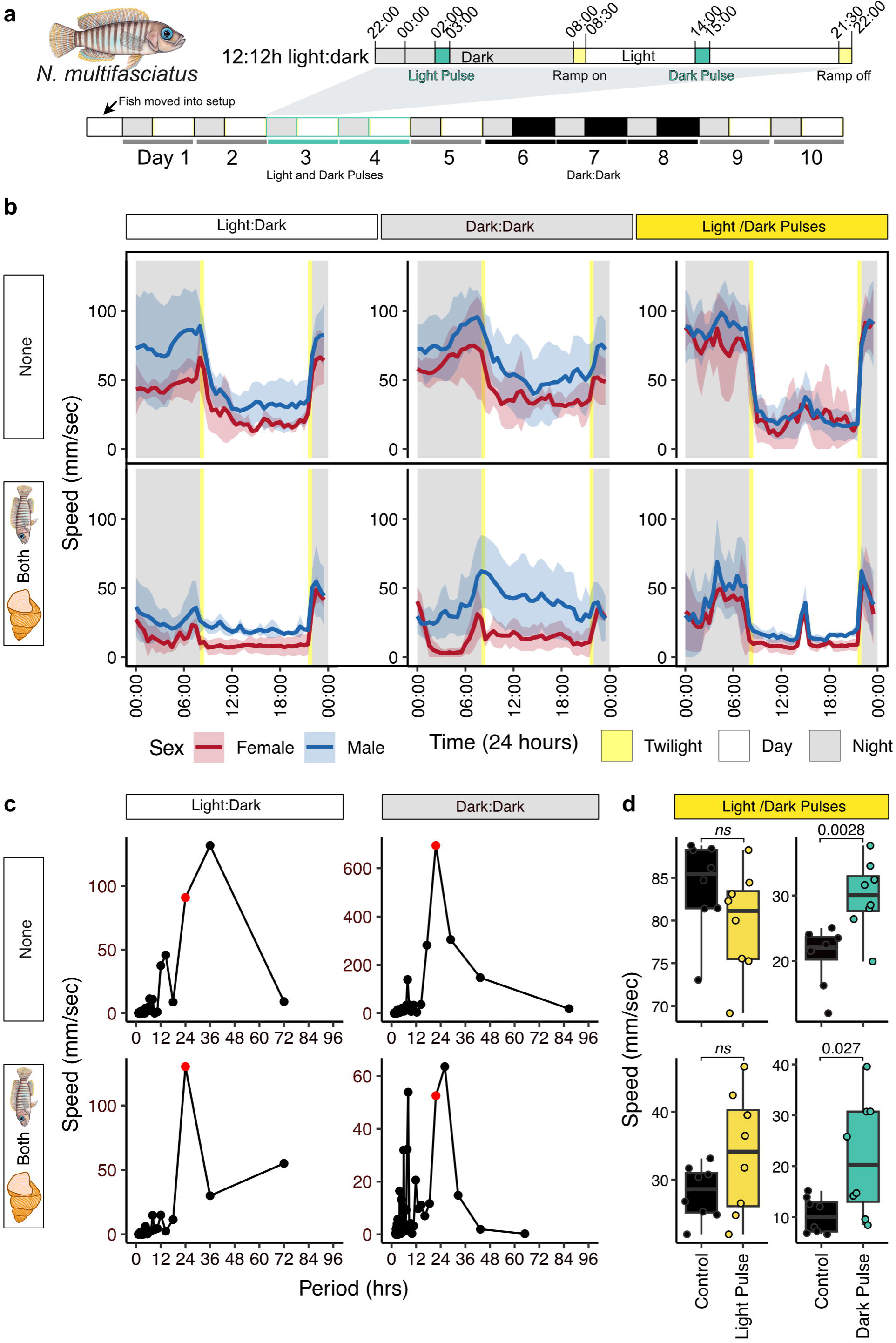
The masking effects of shells and conspecifics are circadian and independent of light. (**a**) Schematic of light cycle for in lab tracking of activity patterns during light perturbations. Grey areas indicate night when lights are off, yellow areas indicate dawn and dusk when lights are ramping up or down, and white indicate daytimes when lights are on. Dark pulses (lights off) and light pulses (lights on) are indicated by turquoise highlights. (**b**) Plots showing the average daily activity patterns for laboratory strain *N. multifasciatus* without any enrichment (top panel) or in the presence of both a shell and conspecifics (bottom panel), during Light:Dark, Dark:Dark, and light pulse periods. Each plot shows the activity patterns of 8-10 individuals, averaged across days. Shaded ribbons display 1 standard deviation from the mean. Grey shaded areas indicate night-time or the dark periods during the day in (B, E), yellow (in A, C, D, F) and dark grey (in B, E) are dawn and dusk, white areas are daytime, green areas are the dark pulses and orange areas are the light pulses. Boxplots show the interquartile range (limits of boxes correspond to 1st and 3rd quartiles), whiskers represent 1.5*(interquartile range), and the line is the mean.

We next hypothesised that if light serves as a masking factor, activity should immediately increase during the day when a dark pulse occurs (lights turned off for 1 hour) and respectively decrease during the light pulse at night (lights turned on for 1 hour). Individuals in both conditions showed identical responses to these manipulations (**Figure 5d**). *N. multifasciatus* individuals displayed a significant increase in activity during the dark pulse compared to the two hours before the pulse, but no corresponding decrease in swimming speed during the light pulse, compared to the two hours before the pulse (**Figure 5d-e)**. These experiments demonstrated that the activity patterns of *N. multifasciatus* are primarily masked by the presence of shells and conspecifics. The response to light pulses may reflect a startle reaction, but it does not appear to alter the underlying activity pattern.

## Discussion

How social and environmental factors influence circadian activity patterns in animals is poorly understood. Here, using species from the model system of Lake Tanganyikan cichlid fishes, we demonstrate roles for conspecifics and physical environmental cues related to their life history traits in shaping temporal activity patterns. Specifically, we demonstrate that conspecific social interaction and empty snail shells act as masking factors to generate diurnal activity patterns in shell-dwelling cichlids. We show that the effect of these cues is species-specific, affected by life history traits and modulated by sex and level of domestication, but is unaffected by light cycle. Social cues were necessary to mask activity rhythms in a highly social cichlid, *N. multifasciatus*, but not the solitary *N. brevis*. Furthermore, these cues are highly salient, and a visual cue of a shell alone was sufficient to elicit a masking effect in female *N. brevis* in the absence of physical interaction. These results extend previous studies on the activity patterns of fish and cichlids from Lakes Tanganyika and Malawi demonstrating how temporal niche partitioning may have facilitated their adaptive radiation (Lloyd et al., 2024, 2021; Nichols et al., 2025; Shafer et al., 2023). Our work suggests that masking of endogenous circadian rhythms could contribute to the generation of diverse activity patterns in these fishes, and supports social interactions as a potent regulator of circadian behaviours across animals.

A masking factor has an immediate and reversible effect on circadian activity patterns, whereas an entrainment factor acts to set or change the underlying circadian rhythm of an organism over time (Mrosovsky, 1999). The phase-shifts in activity patterns that we observe in shell-dwelling cichlids occurred immediately upon entry into the experimental environment with or without shell and conspecific cues. Additionally, the effect of both cues continued in the absence of light, suggesting that they indeed modulate the output of the circadian clock and are *bona fide* masking factors. Traditionally, masking of circadian activity patterns is best understood in relation to light, which has a negative masking effect on activity patterns in nocturnal species, and a positive masking effect in diurnal species (Mrosovsky, 1999). Several other cues have been identified as masking factors, including temperature (Bacigalupe et al., 2003; Halle, 1995), predation, wheel-running (Kas and Edgar, 1999), and, in the case of spiny mice, their environment (field vs laboratory) (Levy et al., 2007). However, our study is the first to identify both conspecific cues and a naturally occurring and physical abiotic cue (empty snail shells) as masking factors for circadian activity. In Mongolian gerbils (Weinert et al., 2007) and degus (Kas and Edgar, 1999) the presence of an artificial running wheel can positively mask activity patterns. However, shell dwelling cichlids have co-evolved with *N. tanganyicense* snails, whose empty shells are widespread and long-lasting throughout Lake Tanganyika (RYAN et al., 2020). *N. tanganyicense* are endemic to Lake Tanganyika, predating its existence(Sengupta et al., 2009; Van Damme and Pickford, 1998), and usage of their empty shells for breeding and shelter by cichlids has most likely evolved multiple independent times (Lein and Jordan, 2021; Ronco et al., 2021). This extensive history could explain the potency of shells as a cue, including the observation that female *N. brevis* were masked by the sight of a shell alone.

The relationship between chronobiology and sociality is complex (Castillo-Ruiz et al., 2012). Interestingly, we observed that the sociality of a species dictates whether conspecific cues have a masking effect. Sociality exists as a spectrum across species of shell dwelling cichlids, with some species remaining solitary except during mating (e.g. *N. brevis*), some which form harems of 3-4 individuals (e.g. *L. ocellatus*), and some forming complex social hierarchies of up to 30 individuals (e.g. *N. multifasciatus*) (Jordan et al., 2021). Our results suggest that masking factors have co-evolved with life history traits in these species. Additionally, consistent with the observed effect of sex on activity patterns and responses to cues, large sexual dimorphisms in the morphology, behaviour, and the relationships individuals have with shells also exist within specific shell-dwelling cichlid species (Schütz and Taborsky, 2000). Integration of social cues and circadian rhythms has also been observed in the Emperor cichlid, which exhibits circadian yawns in males and contagious yawns in the presence of conspecifics (Abdalla-Wyse et al., 2026). Comparisons across species with and without sexual dimorphisms (Taborsky, 2001), species along the sociality spectrum (Jordan et al., 2021; Lein and Jordan, 2021), and species from clades representing independent evolution of shell-dwelling (Ronco et al., 2021) could clarify the direction (positive or negative) and evolutionary origins of the observed masking effects.

Intriguingly, our results suggest that in the absence of environmental and social cues, shell-dwelling cichlids from Lake Tanganyika exhibit nocturnal activity patterns. Previous studies have suggested that the presence of behavioural masking could represent an early step in an evolutionary transition between nocturnal and diurnal activity patterns (Hut et al., 2012). Indeed, such a mechanism could facilitate transitions, and help explain the observed rapid evolution of temporal activity pattern diversity in cichlid fishes (Lloyd et al., 2021; Nichols et al., 2025). However, from our study, it is unclear which activity pattern (diurnal or nocturnal) is ancestral, or if these observations represent positive or negative masking of locomotor activity. For example, the presence of each cue could be a positive mask on diurnal activity, or a negative mask on nocturnal activity (Mrosovsky, 1999). The use of empty snail shells has likely evolved multiple times within the Lake Tanganyikan cichlid radiation from a non-shell using ancestral species (Lein and Jordan, 2021; Ronco et al., 2020). This suggests that shell dwelling species were ancestrally nocturnal, and have evolved a positive diurnal masking effect in response to shells, and representing a potential first step in the evolution of a diurnal lifestyle. However, our previous work has identified many transitions between activity patterns within the Lake Tagnayikan cichlid radiation, most of which have occurred in lineages without the life history trait of shell use (Nichols et al., 2025). Alternatively, masking of activity patterns in shell dwelling species may be an adaptive plasticity (Helm et al., 2017; Kronfeld-Schor and Dayan, 2003), allowing them to be active during night or day under different prevailing environmental conditions. Again, additional information from other shell dwelling species, as well as a more complete phylogenetic coverage of activity patterns could help distinguish between these different hypotheses and allow a more accurate reconstruction of ancestral patterns of circadian activity within this clade.

Importantly, the masking effects of shells and conspecifics were not observed for wild-caught fish that were recently brought into the laboratory setting. This suggests the existence of additional masking factor(s), which were either lost from the lab and aquarium strains during domestication, or imparted by experience in the wild. In a similar paradigm, the well studied Golden spiny mouse is known to display diurnal activity patterns in their natural habitat, but convert to nocturnal activity patterns when isolated in a laboratory setting (Cohen and Kronfeld-Schor, 2006; Levy et al., 2007). This switch in activity has been primarily attributed to the presence of intra-specific competition with the nocturnal *A. cahirinus (Kronfeld et al., 2000)*, forcing the spiny mice into the diurnal niche, and representing a switch from a nocturnal ancestor (Levy et al., 2007). Under an analogous scenario, shell-dwelling cichlids might be ancestrally or defaultly nocturnal, but forced into diurnal niches in the lake due to competition with other shell-dwelling cichlids, or predation pressure from catfishes or other nocturnal piscavores (Peart et al., 2014). The presence of conspecifics in social species, as well as empty snail shells, could provide additional benefits to a diurnal lifestyle, including shelter and “safety in numbers” from predators at night, allowing shell dwellers to rest during the night when they would otherwise be unable. For domesticated strains of *N. multifasciatus*, they may either have lost the masking effect due to the absence of competition and predation, or never learned that association in the aquarium. Alternatively, there could be unaccounted for abiotic factors (temperature, pH, photoperiod), or biotic factors such as specific food sources, or non-predatory intra-specific interactions that influence activity in wild caught fish. In the original definition of a masking factor, Fry (1947) (Fry, 1947) suggests that they may be far-reaching, defining them as any “factor which prevents a second identity from operating on the organism to the extent that it would if the masking factor were not present,” and that masking factors might be so ubiquitous that they “would appear rarely to operate per se.” Our results suggest that multiple factors, some known, and some unknown, work in concert to control and mask the activity patterns of shell dwelling cichlids. Indeed, it is possible that masking may be an underappreciated, yet widespread phenomena controlling temporal activity patterns and their evolution across species.

## Methods

### Monitoring of diel activity patterns in the wild

To investigate the diel activity patterns of shell-dwelling cichlids in their natural environment, we designed an infrared camera system to record 24-hour video sequences *in situ* in Lake Tanganyika. We examined three species in the southern part of Lake Tanganyika. Populations of the focal species *N. multifasciatus* were investigated at Chikonde Bay, Lake Tanganyika, Zambia (8°42’49.4” S, 31°07’23.0” E), and *Lamprologus ocellatus* and *Telmatochromis temporali*s were examined at Kalambo Lodge, Lake Tanganyika, Zambia (8°37’25.3” S, 31°12’3.4” E). The camera system was mounted approximately 50 cm above shell beds to ensure optimal image quality while minimizing anthropogenic disturbance, and territories for recording were randomly selected each day. The system was demounted and reinstalled daily before noon. All recordings were done in Fall 2022. To measure overall activity of each species, fish were manually tracked from each video recording, and their individual movements counted, by summing the number of times fish appeared in the frame for the full length of the video. This method provides a measure of overall activity, but does not accurately enumerate the number of individuals at each location. Dawn/dusk times were taken from https://www.timeanddate.com/sun/zambia/lusaka?month=10&year=2022. All experiments in Zambia were performed under approved study permits.

### Infrared field camera

Field observations were conducted using a custom-built setup consisting of modified Hero 3 GoPro cameras with 6 mm CCTV M12*0.5 IR lenses, controlled by an integrated Raspberry Pi system (Raspberry Pi Zero W with a Waveshare Relais Board) and enclosed in an aluminum housing. The IR-blocking filters of each Hero 3 GoPro camera were removed. The Raspberry Pi simultaneously controlled the infrared (IR) illumination using 940 nm LED lights (intensity: 19.2 W/m²; 7.68 W used in the setup). Recordings were conducted for 10 minutes every hour across a roughly 24-hour period. The system was powered by a Samsung INR18650-35E battery pack and mounted on a tripod that could also be arranged as a four-leg aluminum frame (diameter = 30 mm).

### Animal husbandry

We used three different strains of *N. multifasciatus* and one strain of *N. brevis*. All fish were housed in the fish facility of the Biozentrum, University of Basel, Switzerland. The three *N. multifasciatus* strains differed in the number of generations they had spent in captivity and in their rearing conditions. The aquarium strain (AS) was obtained from a commercial breeder (Cichlidenstadl, Blindheim, Germany) in Spring 2022 and was kept in captivity for an unknown number of generations. The wild strain (LS) was originally imported in 2018 and was kept in captivity for approximately 2-3 generations, and housed in conditions consistent with their natural environment (in social groups and with natural or 3D printed shells). Finally, the Zambian strain (ZP) was imported in October 2022, and was held under laboratory conditions for less than two months before experimentation under the same conditions as the LS. *N. brevis* individuals were obtained from the Max Planck Institute of Animal Behavior, Konstanz, where they were housed in conditions similar to their natural environment for one generation. All fish were kept on a 14 h:10 h light–dark cycle (lights on 08:00–22:00, lights off 22:00–08:00). During the experiment, all fish were fed every second day in the morning with dry food flakes (Dr. Bassleer biofish-food (Aquarium Münster)) and with live artemia. All animal experiments were approved by cantonal authorities (licence #3102, Canton of Basel-Stadt, Switzerland).

### 3D printing artificial shells

Artificial shells were chosen to standardize husbandry conditions. Furthermore, because the cichlids hide inside the shells, the use of snap-fit closures allowed the shells to be opened easily, reducing stress during handling and transfers(Bose et al., 2020b). We avoided the use of magnets, glue, and metal parts in the design of the snap fit closing, as prototypes with these materials were prone to degradation of the glue and rusting. We modified an available 3D model of a *Neothauma tanganyicense* shell(Bose et al., 2020b) and equipped it with a snap fit closing (using the software Blender 3.1, Fusion360, and Moment of Inspiration). We then 3D-printed this model using an SLA printer (Form 3/Form 3L, Formlabs) with a layer thickness of 0.100 mm using white resin. After printing, the shells were immersed in an isopropanol bath to remove the support structures, then dried and UV-cured (Form Cure/Form Cure L, Formlabs).

### Diel behaviour tracking

We tracked fish using a previously published protocol and software(Nichols et al., 2024). Briefly, fish were recorded from the front of each tank with Spinnaker Chameleon 3 monochrome CCD cameras (model CM3-U3-13S2M-CS RoHS 1.3MP B&W Chameleon USB 3.0 Camera 1/3” CCD CS-Mount) (FLIR). LED infrared (IR) light panels illuminated the tanks from the back, while overhead white lights provided daytime, dawn, and dusk illumination. Cameras were equipped with Fujinon YV4.3x2.8SA-2 CS-Mount 2.8-12mm Varifocal lenses capped with long-pass filters that permitted only infrared light (IR) to be detected by the cameras. The experimental fish were tracked at 10 frames per second (fps) with a custom written code in Python 3.7 using the PySpin module to control cameras. Fish positions were extracted from each frame, and the position of each fish was used to calculate the speed over time (mm/s). Fish tracks were then analysed using custom code written in R. All code is available online (Python scripts: https://github.com/maxshafer/cichlid-analysis-mers; R code: https://github.com/maxshafer/R_tracking_analysis).

### Experimental manipulation of social and environmental cues

In total, we examined 120 adult individuals of *N. multifasciatus* and 50 adult individuals of *N. brevis* in an equal sex ratio. Our experimental setup allowed the testing of 6-10 individuals per session as previously(Nichols et al., 2025). Specifically, individual fish were alternatively placed in every second arena and separated with mesh dividers (**Supplemental Figure 1A**). On the day the fish were transferred into the experimental tank, recordings and real-time tracking started at midnight, following an acclimation period of approximately 12 hrs. Each fish was recorded for 6 days and 6 nights (144 hours). For each strain of *N. multifasciatus* (AS, LS, and ZP) and *N. brevis*, we tested a minimum of 4-6 individuals of each sex under four different experimental conditions; (*i*) in the absence of all cues (no shell, no conspecifics), (*ii*) in the presence of a single shell, (*iii*) in the presence of conspecifics, and (*iv*) in the presence of both a single shell and conspecifics. For experiments in which individual fish were tested in the presence of conspecifics, we first assigned them to groups of three (one male and two females) one week prior to the start of the recording to establish familiarity. For the experiment, the groups were separated: one focal fish was placed in an arena (with or without a shell), while the remaining two fish were placed in an adjacent arena with shells. The focal fish and its associated social group were separated by a mesh divider, allowing visual interactions. To obtain 4-6 individuals of each sex, group fish were reused to randomly establish new groups for subsequent experiments, and focal fish were removed from further testing.

### Light cycle manipulations

To investigate the masking response of shells and conspecifics on *N. multifasciatus* to different light conditions we followed the above protocol but with several modifications. Light perturbation experiments lasted 10 days and 10 nights (240 hours). Light and dark pulses were performed by switching the lights on or off for 1 hr during the nighttime and daytime, respectively, and fish were subsequently exposed to three days of continuous darkness (dark:dark). Light perturbations (pulses, dark:dark) were separated by 48 hours of ‘normal’ light cycle periods with 14 hours light and 10 hours dark. Fish were exposed to ‘normal’ light cycle periods for 48 hours at the beginning and again after perturbations (**Figure 5a**).

### Analysis of behavioural data, statistical analysis, and plots

Raw fish behavioural tracks were generated by Python code as previously described(Nichols et al., 2025) (**Supplemental data 2-3**). All plots were generated by, and all analyses were performed in R (version 4.5.0). Analysis of variance (ANOVA) was used to determine the contributions of various factors to swimming speed (presence or absence of cues, strain, sex), followed by post-hoc analysis using TukeyHSD or students t-tests to determine specific differences. The fft() function from the default stats package was used to perform Fourier analysis of periodic patterns in swimming speed. All statistical tests were performed in R (version 4.5.0). The R package lubridate was used for manipulation of data with associated dates and times(Grolemund and Wickham, 2011). We used the patchwork package to design and produce plots.

### Data and code availability

Field recordings and processed animal tracks for all in lab experiments are available via dataverse at the University of Toronto (https://doi.org/10.5683/SP4/MNW5G8). All code is available on github with separate repositories for tracking software (https://github.com/annnic/cichlid-analysis) and R analysis (https://github.com/maxshafer/R_tracking_analysis).

## Acknowledgements

We thank members of the Shafer, Salzburger, and Schier labs for helpful advice and feedback. We thank Alex Schier for helpful advice and feedback on project design and use of aquatics facilities. We thank Dani Lüscher, Sven Gfeller and Simon Saner for help with the design and construction of the field camera system. We thank both Richard Michael Newton and Thomas Garello of the Research Instrumentation Facility of the Biozentrum Basel for their support and assistance in the design and printing process of the artificial shells. We thank Alex Jordan for providing the original model of the shell, as well as the laboratory strain of *N. brevis*, and advice regarding locations of specific species in the field. We thank Julie Johnson for the cichlid illustrations. This work was made possible by logistical support from divers and staff at Kalambo Lodge, Zambia. This work was supported by grants from the Natural Sciences and Engineering Research Council of Canada (NSERC) (RGPIN-2024-05509) to M.E.R.S., and the Swiss National Science Foundation (SNSF) to M.E.R.S. (196313) and W.S. (208002).

## Author contributions

A.L.A.N., W.S., and M.E.R.S. conceived and designed the study. L.F., A.L.A.N., and M.E.R.S collected behavioural data. L.F. and M.E.R.S. analysed behavioural data. R.G.D., A.I., and A.R. aided in study design and collection and interpretation of behavioural data. L.F., W.S., and M.E.R.S. interpreted the results and wrote the manuscript. All authors read and approved the manuscript.

## Notes

### Competing Interest Statement

The authors have declared no competing interest.

